# Evaluation of the effect of life extension of DHEA extract from sweet potato on *Caenorhabditis elegans*

**DOI:** 10.1101/2020.12.16.423044

**Authors:** Huan Chen, Dan Zou, Lufeng Wang, Junyi Huang

**Affiliations:** College of Food Science and Technology, Huazhong Agricultural University, Wuhan, Hubei, 430070, China; Key Laboratory of Environment Correlative Dietology, Ministry of Education, Huazhong Agricultural University, Wuhan, Hubei, 430070, China; School of Life Sciences, Shanghai University, NO. 381 Nanchen Road, Shanghai 200444, China

**Keywords:** Sweet potato, Dehydroepiandrosterone, *Caenorhabditis elegans*, antiaging

## Abstract

Products from natural sources are being used from centuries. This study investigates the potential antiaging activity of DHEA extracts from sweet potato. DHEA was extracted with the assistance of acid hydrolysis-ultrasonic, and the *Caenorhabditis elegans* model was used to investigate the antiaging activity. The results from the current study articulated that DHEA from sweet potato in 50 μmol / L effectively prolonged the life-span of *C. elegans* by 13.60%, but the effect was not concentration-dependent. Besides, DHEA had no effect on the growth of *E. coli* OP_50_ and the nematode pharyngeal pump rate, indicating that DHEA didn’t delay the aging of the nematode through calorie restriction. Further experiments demonstrated that DHEA would slow the growth of nematode body size, reduce the accumulation of ROS and lipofuscin of *C. elegans*. The expression and migration of *daf-16* into the nucleus of nematode were significantly improved as well. The antiaging effect of DHEA on *C. elegans* may be achieved by strengthening the nematode’s ability to resist oxidative stress and promoting nuclear expression of the *daf-16* gene.

**Summary Statement:** This study proved that DHEA from sweet potato can extended lifespan of *Caenorhabditis elegans* induced by strengthening the antioxidant capacity and promoting the nuclear expression of *daf-16*

## INTRODUCTION

*Ipomoea batatas* Lam, also known as sweet potato, is one of the three major potato crops, which possess high edible value. Sweet potatoes have been cultivated in more than 100 countries because of their capacity to adapt to various soils, strong resistance to stress, and good cold and barren tolerance, Furthermore, sweet potatoes play a significant role in laxatives, anti-diabetes, antiaging, etc.(Loening-Baucke et al., 2004; Mohanraj and Sivasankar, 2014). Studies have shown that sweet potatoes contain a variety of physiological substances with numerous biological activities, and Dehydroepiandrosterone (DHEA) may be one of the most influential (Ran, Liang, Du, & Sun, 2019).

Dehydroepiandrosterone is a precursor of sex hormones, a steroidal compound secreted by the human adrenal cortex, which plays a very important role in the synthesis and secretion of other hormones by the adrenal gland(Genazzani et al., 2007). DHEA can be converted into estrogen or androgen by enzyme catalysis in circulating blood. Some scholars used wood residue rich in plant sterols as raw materials to synthesize DHEA in 1953, but the product yield was too low, the synthesis process was cumbersome and not conducive to environmental protection. In order to address these problems, using esters of diosgenin to synthesize DHEA has been discovered in 1957, which is the more common method to produce DHEA in the modern industry. Moreover, Stephanie has successfully synthesized DHEA and other steroids through biosynthesis in 2006 (Webb et al., 2006). However, it must be cautious that the physiological activity of synthetic DHEA may be significantly different from that of natural DHEA, so the extraction of natural DHEA is particularly a more significant contribution to science. Hu Gunnel successfully attained DHEA from yam in 2005, and Ran has greatly improved the extraction rate of DHEA in sweet potatoes pomace by ultrasonic-microwave synergistic employing(Nordmark et al., 2005) (Ran et al., 2019).

Many investigations have demonstrated that DHEA as a sex precursor has a wide range of biological activities such as regulating obesity, regulating blood sugar levels, enhancing resistance, anti-cancer potential, anti-viral activity, and relieving stress(Barkhausen et al., 2009; Barrettconnor and Edelstein, 1994; Deleon et al., 2002; Webb et al., 2006). The literature of Grillon has revealed that DHEA also has the physiological effect of delaying aging(Grillon et al., 2006).

The current investigation was done to clarify the antiaging effect of DHEA and explore the health value of sweet potatoes. In this study, sweet potatoes were as raw materials to extract and purify DHEA. And the life expectancy, physiological and biochemical indicators of *Caenorhabditis elegans* served as evaluation indicators to establish an aging model to examine the theoretical basis of sweet potato antiaging. *Caenorhabditis elegans* is a non-toxic, independent living organism with high safety. The results derived from these experiments can be directly applied to higher organisms, even humans owing to the similarity in tissue structure and life-span regulation mechanism between them(Wang and Xing, 2010a).

## MATERIALS AND METHODS

### Materials

Sweet potatoes were purchased from the market named Baishaizhou (Wuhan China). Wild-type N2 *Caenorhabditis elegans* were obtained from the Caenorhabditis Genetics Center. Uracil-deficient *E. coli* OP_50_ was kept by the laboratory. Fluorouracil (FUDR) was provided by Sigma-Aldrich Inc. DHEA and other chemicals were purchased from Aladdin Reagent Co., Ltd (Shanghai, China). All of the materials used in the study were all of the analytical grade.

### Preparation of DHEA

DHEA was extracted by acid hydrolysis-ultrasonic assisted following method of Ran with slight modifications(Rangsinth et al., 2019). The sweet potatoes were washed and beaten into fine particles. And the particles were cultivated at 60 °C for 24 h. Then, 3% concentrated sulfuric of fresh sweet potato was added, and the stable suspension was hydrolyzed in the water base of 90 °C for 36 h. After the suspension cooling to room temperature, adjusted its pH to 7. Following the suspension was washed 2 to 4 times with distilled water and was centrifugated in a high-speed refrigerated centrifuge for 2 min at 10000 g. Separated the remaining residue and freeze-dried it to obtain a solid powder sample. Then, the collected sample was put into a microwave oven for 40 min at 0.5 W / cm^2^, 40 kHz to extract DHEA. Rotate the extract and collect it. Transferred the extract to a silica gel column (ether: acetone = 1: 1) to remove non-polar substances. The elution was gathered to dryness in a vacuum oven to yield a solid extract. Prepared DHEA samples (28.8 mg, 21.6 mg, 14.4 mg, 7.2 mg, and 0 mg) were precisely weighed and dissolved in 1 mL of dimethyl sulfoxide (DMSO) respectively. Therefore, a series of DHEA standard solutions (100 mmol / L, 75 mmol / L, 50 mmol / L, 25 mmol / L, 0 mmol / L) were obtained. Samples were kept the freezer at −20 °C.

### Culture of *C. elegans*

When *C. elegans* grew into an adult, they were cultivated on the plate of NGM medium freshly coated with *E. coil* OP_50_ for 2-4 h to lay eggs. NGM plates with eggs were cultured in a constant temperature incubator at 20 °C to acquire age-synchronic nematodes(Peixoto et al., 2019; Wang et al., 2020). The nematode growth broth was ready as following.

S. Medium I: S basal 100 mL, 1 mol / L potassium citrate solution (pH 6.0) 1 mL, Trace Metal Solution 1 mL, 1 mol / L CaCl_2_ 100 μL, 1 mol / L MgSO_4_ 100 μL, 5 mg / mL cholesterol solution 100 μL was mixed up in the sterile operating platform. S. Medium II: FUDR solution and *E. coli* OP_50_ were added to S. Medium to ensure their concentrations 150 μmol / L and 5 mg / mL, respectively.

### The life-span assay of *C. elegans*

The synchronized nematodes that in satisfactory growth state under sterile conditions were picked for this assay. *C. elegans* were divided into experimental groups treated with 1.1 μL various concentrations of DHEA (0 mmol / L, 25 mmol / L, 50 mmol / L, 75 mmol / L, 100 mmol / L), and the control group with 1.1 μL S. Medium I. All of them were placed into EP tubes with 1.1 mL S. Medium II. After mixed each of them, injected them into a 96-well plate, 100 μL per well, a group of 10 wells and 10 nematodes a well, respectively. Then cultivated them in a constant temperature incubator at 20°C. The day when nematodes were picked was regarded as 0 days of their life. Observed the nematode status and recorded the number of nematodes in each group daily until the nematodes were all dead. The loss and the death of worms were not counted in the statistical data. The experiment was repeated three times.

### Bacterial growth assay

*E. coli* OP_50_ is the food for wild-type *C. elegans* N2, which was utilized in this assay. The experiment was conducted in a 96-well plate. Selected 28 wells and divided them evenly into 7 groups (0 mmol / L, 25 mmol / L, 50 mmol / L, 75 mmol / L, 100 mmol / L, S. Medium, LB liquid medium). Each well of the plate contained 200 μL LB liquid medium containing *E. coli* OP_50_ and 0.2 μL of the corresponding concentration of DHEA solution. The 96-well plate was later placed in a constant temperature incubator at 20°C. Measured OD_600_ of each well every 1 hour, and recorded a total of 8 hours. The assay of the effect of DHEA on *E. coli* OP_50_ was performed in triplicate.

### Antioxidant stress assay

Selected synchronized nematodes in good condition and put them into 12-well plates, and divided the well into 6 groups, a group with 10 wells and a well of 10 worms. Treated the worms with 90 μL mixed liquid of which 1 μ L DHEA solutions (0 mmol / L, 25 mmol / L, 50 mmol / L, 75 mmol / L, 100 mmol / L) and 1 mL S. Medium II or 1μL S. Medium I and 1 mL S. Medium II. Added 10 μL of 10 mmol / L H_2_O_2_ solution to each well under light-proof conditions after incubating the nematodes for 6 days under 20°C. Subsequently, they were cultured in a seal and dark constant temperature incubator at 20°C. Monitored the nematode status every 1-2 hours and counted the number of nematodes that remained in each group. But the loss or death was not involved in the statistical data. All tests were carried out for a total of three times.

### Measurement of body size

*C. elegans* were cultivated and treated with or without the DHEA solutions from sweet potatoes as in the life-span assay described above. The day when the nematodes first treated with DHEA was considered as day 0. Picked one of the nematodes in good condition in each well to take photos on the 3rd, 6th and 9th days under utilization of Leica Application suite V3 software (version 3.40). Then used Photoshop software to measure the length and width of the nematode. All determinations were replicated three times.

### Pharyngeal pumping rate assay

The worms were cultivated on the 96-well plates with various concentrations of DHEA or S. Medium as the control group, as mentioned in the life-span assay. The microscope and Leica Application Suite V3 software (version 3.40) was applied to determine the pharyngeal pumping rate every 20s of *C. elegans* on the 3rd, 6th and 9th days. The experiment was repeated three times.

### Locomotion Assay

The synchronized nematodes were raised on 96-well plates similar to the life-span assay. Briefly, the wells were divided into 6 groups evenly, and 200 μL of mixed liquid, which was made by 0.5 μL of a series of DHEA solution and 0.5 mL S. Medium II (experimental group) or 0.5 μL S. Medium I and 0.5 mL S. Medium II (control group) was transferred to each well. The frequency of body bends of nematodes was recorded on day 3, day 6 and 9. Worms treated with or without DHEA was picked up to measure the body bends rate in 30 s with a stereo microscope. And the bend of *C. elegans* was defined as the process of taking a S-shaped body formed as the central axis as the axis and traveling a sinusoidal waveform forward(Ruan et al., 2016; Wang and Xing, 2010b). The assays were performed in triplicate.

### Accumulation of ROS in *C. elegans*

2,7-dichlorodihydrofluorescein diacetate (H_2_DCFDA) was served to detect ROS levels in nematode cells. First, synchronized L4 larvae were maintained in DHEA solutions as described above for 7 days. Then, worms were washed with 0.1% PBST (1 × PBS / 0.1% Tween 20) buffer to wipe off the *E. coli*. Subsequently, 10 washed worms were placed in 80 μL of PBS in a well of a 96-well blackboard. At the same time, 20 μL H_2_DCFDA(250 μmol/L)was transferred to the well to ensure the final concentration of H_2_DCFDA in the well is 50 μmol / L(Pant et al., 2014). Fluorescence was conducted at excitation/emission wavelengths of 485 nm/535 nm every 20 min for 8 h at 37°C by using an Agilent chnologies microplate reader. The time point of maximum absorption value was selected for calculation of the DHEA’s inhibition rate of ROS in *C. elegans*. The assay included blank group (without nematode and H_2_DCFDA) and background group (with nematode only and without H_2_DCFDA). And the assay had to be carried out three times.

### Measurement of Lipofuscin

Synchronized nematodes at Larva 4 stage were chosen for this assay. *C. elegans* were retained and treated similar to the life-span assay. The nematodes on the 10th day of synchronism was attached on a 2% sodium azide anesthetized glass slide to a 3% agarose pad and covered with a cover slip. A DMI 3000 fluorescence microscopes (× 10) was used to take microphotographs with DAPI filters (excitation wavelength 340-380 nm, emission wavelength 435-485 nm). And lipofuscin fluorescence of the samples was measured quantified with software ImageJ (NIH). All experiments were independently repeated three times.

### Nuclear location assay of *daf-16*

The nematode TJ356 (*daf-16* :: GFP) used in this assay would fluorescently label *daf-16*, which was the key gene of the downstream of the insulin signaling pathway. Diverted the 2nd old Synchronized worms to the NGM plates with or without DHEA for 2 h. Subsequently, the worms were placed on a microscope slide coated with 2% agarose and 1% sodium azide. Detected the distribution of fluorescent protein under a fluorescence microscope (excitation wavelength: 460-495 nm; emission wavelength: 510-550 nm; Nikon-eclipse 80i) at 100-fold magnification to determine the nuclear expression.

### Statistical analysis

All data was analyzed by GraphPad Prism 7.0 and IBM SPSS Statistic 21.0, and expressed as mean ± SD. One-way analysis of Variance (ANOVA) was employed for comparing the difference between groups. The Long-rank (Mantel-Cox) Test was for the analysis of significant difference test. The symbol “*” means the level of statistically significant was at p < 0.05, “**” means the level of statistically significant was at p < 0.01.

## RESULTS

### DHEA prolonged the life-span of *C. elegans*

Life-span is the most direct and important index to measure the aging degree of *C. elegans*. As Table 1 shown, under the suitable environment, the mean life-span of nematodes in the blank control group was 18.5 ± 3.8 days. The survival time of nematodes treated with 50 μmol/L DHEA was significantly prolonged with the survival rate extended by 13.6%. However, there was no significant difference in the mean life of *C. elegans* among the blank control group and the 25 μmol/L, 75 μmol/L and 100 μmol/L groups (P > 0.05), which indicated that the effect of DHEA on delaying nematode aging is not concentration-dependent.

**Table 1.** Life-span of *C. elegans* treated with different concentrations of DHEA.

| Group (μmol/L) | Fatality/amount | Mean life-span (d) | Maximum<br>Life-span (d) | Life extension (%) |
| --- | --- | --- | --- | --- |
| 0 (Control) | 49/53 | 18.5 ± 3.8 | 23 | - |
| 25 | 48/51 | 18.2 ± 4.3 | 23 | - |
| 50 | 49/55 | 21.1 ± 3.5* | 24 | 13.6 |
| 75 | 44/52 | 19.0 ± 4.6 | 24 | - |
| 100 | 44/53 | 19.1 ± 4.6 | 24 | - |
| S. Medium | 52/59 | 19.0 ± 3.7 | 24 | - |

“\*” means $P < 0.05$ compared with control group.

### Effect of DHEA on E. coli OP_50_ Growth

Experiments have been carried out that limiting calorie intake is the most effective way to prolong life so many years ago(Viswanathan et al., 2005). *E. coli* OP_50_ is the food of wild-type *C. elegans* N2. As food sources, its concentration has a crucial meaning in the antiaging model of *C. elegans*. The research findings are given in Fig 1. The results illustrated that the OD_600_ values (at 600 nm, the concentration of *E. coli* OP_50_ bacteria is linearly related to its absorbance) of each group were not significantly different for 8 h at 20°C (P> 0.05).

**Fig. 1.**
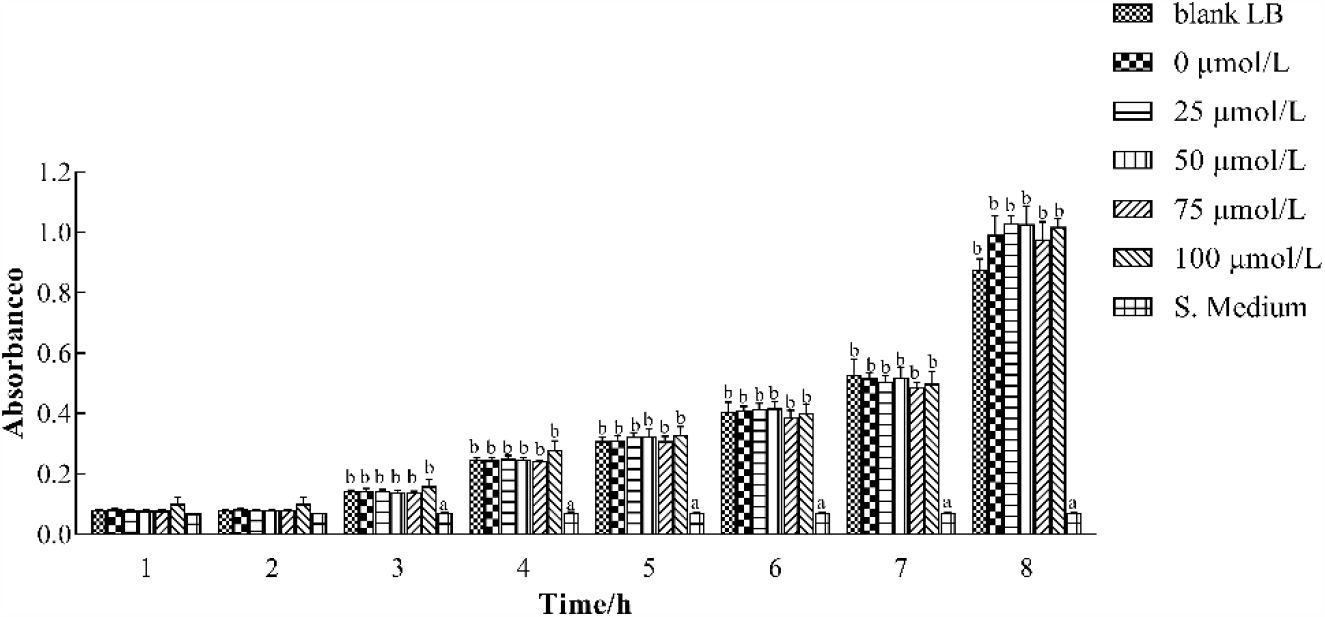
Effect of DHEA on E. coil OP50 growth on 1^st^, 2^rd^, …… 7^th^, 8^th^ hour. Different letters represent significant differences between the groups of each hour.

### Protection against Oxidative Stress

Excess generation of reactive oxygen species (ROS) and reduction of antioxidant defense system capacity will cause the accumulation ROS and the redox reactions in the body, which ultimately result in damage to intracellular proteins, DNA, lipids, sugar water compounds and other substances(Labuschagne and Brenkman, 2013). And it is one of the main reasons for inducing the occurrence of diseases such as cancer, depression, diabetes, Parkinson’s and heart disease. Oxidative stress is a major factor for illness or aging in nematodes-based biological models(Chege and McColl, 2014; Ogawa et al., 2016). The stimulation of paraquat, juglone and hydrogen peroxide will result in oxidative stress damage of *C. elegans*(Chen et al., 2014a; Ku et al., 2013) . In order to test the effect of DHEA against oxidative stress, synchronized worms were exposed to DHEA solutions in different concentrations. As shown in Fig 2, compared with the blank control group, worms treated with 50 μmol / L DHEA had a certain resistance to oxidative damage caused by H_2_O_2_ (P <0.05), and the effect was most prominent in 8-10 h. The 25 μmol / L, 75 μmol / L, 100 μmol / L DHEA groups and S. Medium groups had no significant difference compared with the blank control group (P > 0.05). This was similar to the effect of polysaccharide from the roots of *Lilium davidii* on the life-span of *C. elegans* under oxidative stress (Hui et al., 2020).

**Fig. 2.**
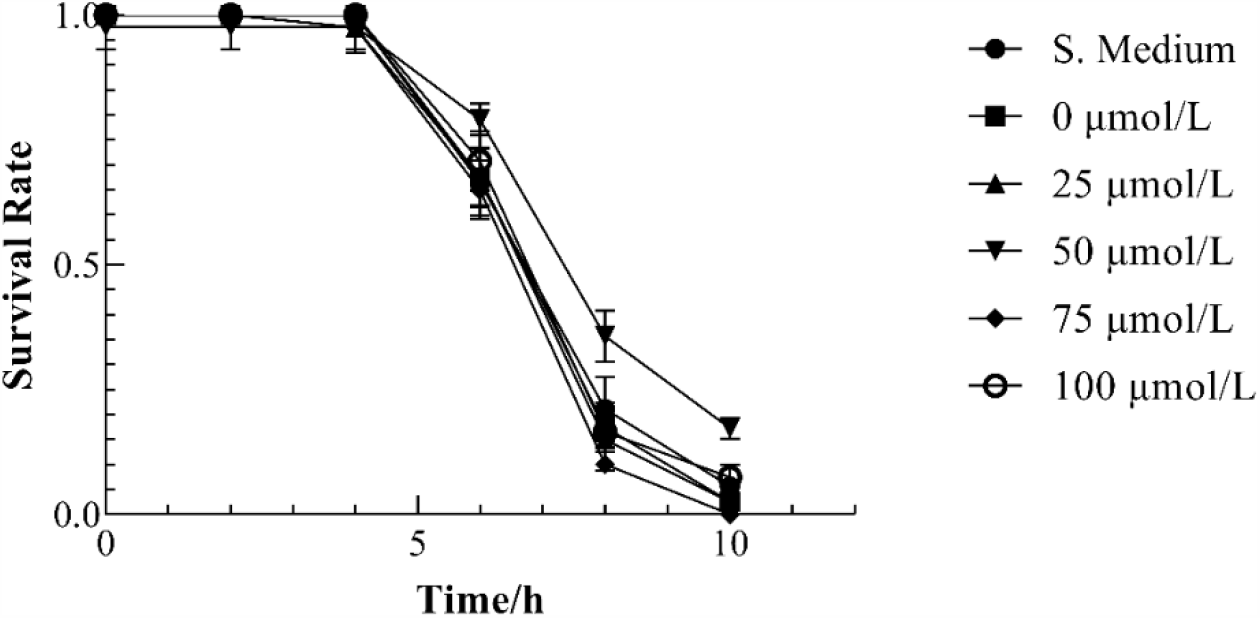
Survival curve of *C. elegans* treated with different concentrations of DHEA.

### Effect of DHEA treatment on body size

Healthy nematodes grown to adult was about 1 mm in length. When nematodes are in a befittingly growing condition, their body length and body width increase gradually in the primary and middle period life. Body length and body width are vital indexes to judge whether nematodes are in normal developmental state. We explored the effect on the body size of *C. elegans*, the results were given in Table 2. The body length and body width of nematodes gradually increased with the growth of age in the primary and middle period of the life of nematodes. Compared with the blank control group, all experimental group inhibited the growth of nematodes and slowed the growth of their body length and body width. There was no significant difference between S. Medium group and the blank control group (P > 0.05).

**Table 2.**
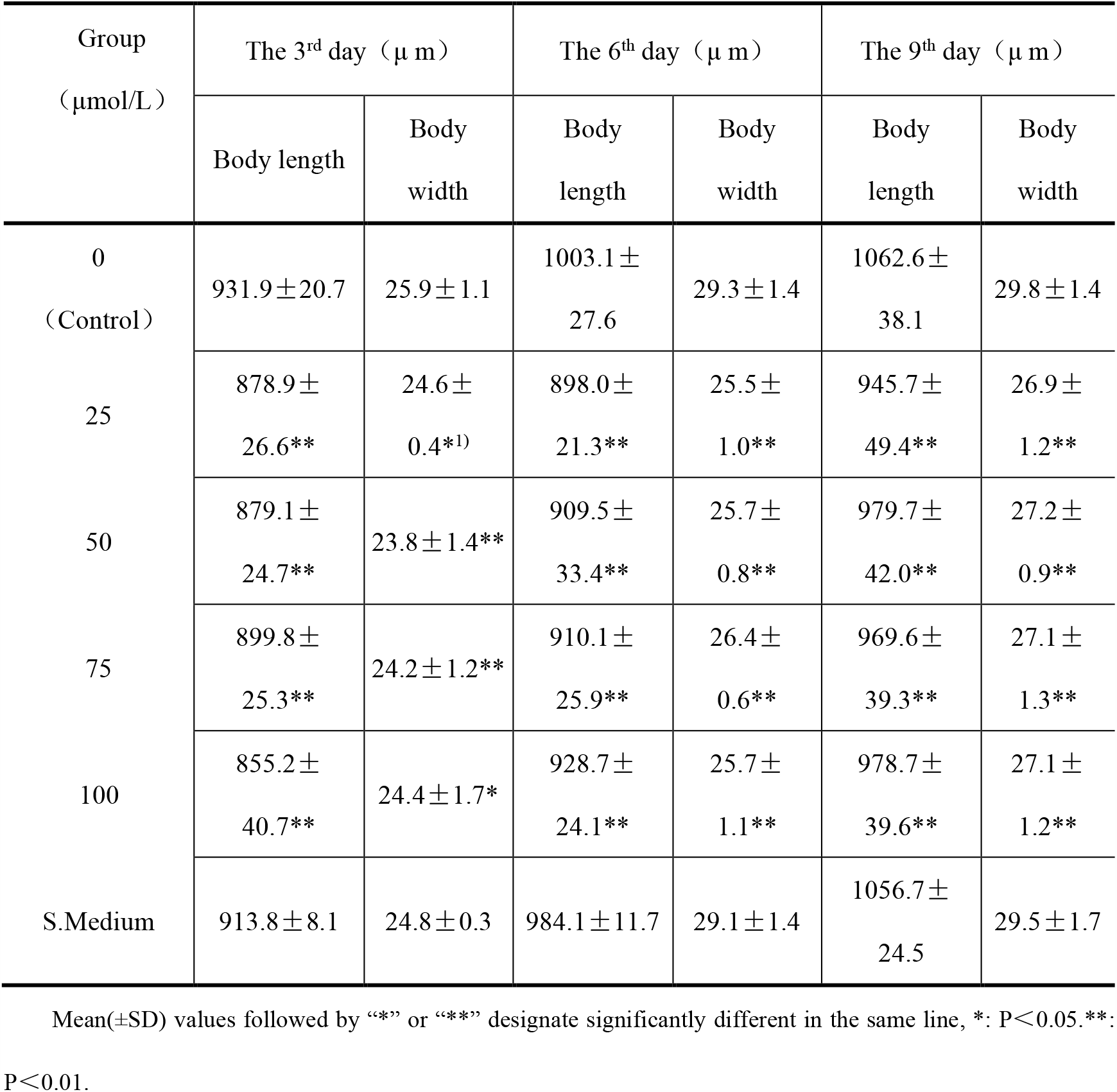
Effect of DHEA on body length and width of *C. elegans*.

### Effect of DHEA on Pharyngeal Pump rate of *C. elegans*

Calorie restriction (CR) has been certified that it can extend the life-span of many species (Ludewig et al., 2004), which is the only known way to prolong the life-span of mammals (Lin et al., 2002). The explanation of CR includes the reduction of cell division, metabolic rate, free radical production (Sohal et al., 1994), and DNA restriction (Masoro, 2005), as well as the stimulating effect of poison (Masoro, 2005; Sohal et al., 1994). The swallowing action of nematodes is correlated with the life-span of nematodes. The normal swallowing frequency of young nematodes is between 150 and 250 times per minute. As age increasing, the nematode’s swallowing action gradually slow down. We can determine the dietary intake of nematodes by monitoring the nematode Pharyngeal Pump rate. Fig 3 illustrates an aging-related decrease, and as we can see, compared with the blank control group, no significant difference was found in 25 μ mol / L, 50 μ mol / L, 75 μ mol / L, 100 μ mol / L DHEA and S. Medium groups (P> 0.05).

**Fig. 3.**
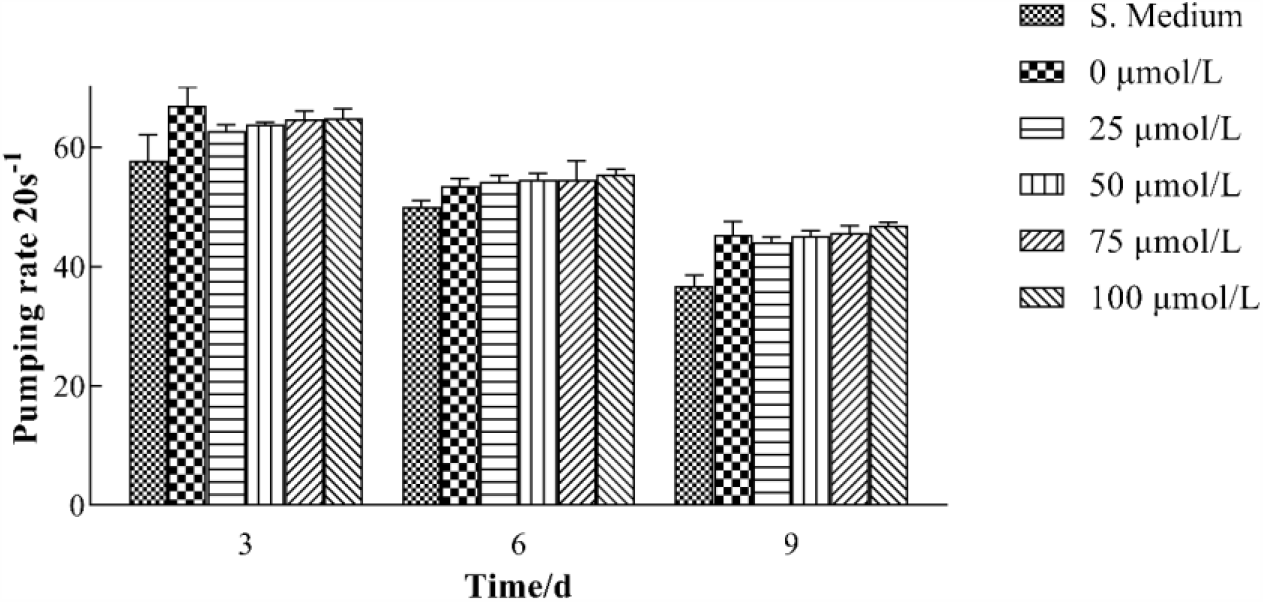
Pumping rate of *C. elegans* treated with different concentrations of DHEA on day-3, day-6, day-9. All groups are not significantly different(P> 0.05).

### DHEA treatment affect the movement of Nematodes

Nematodes in good condition will swing regularly. But as time goes by, the worm’s muscles will contract more slowly, and resulting in a slower rate of swinging or not moving. For the sake of assessing the nematode’s physiological state and aging process, we can evaluate the frequency of nematode movement. As shown in Fig 4, the frequency of swinging of nematodes in each group decreased with age. In the early stage of the nematode life-span (days 3 and 6), the frequency of nematode movement was not significantly different from that in the blank control group (P> 0.05). On day 9, both 25 μmol / L and 50 μmol / L groups increased the movement frequency of the nematode (P <0.01), while 75 μmol / L group has no significant difference (P > 0.05), the 100 μmol / L group slowed down the frequency of nematode movement (P <0.05).

**Fig. 4.**
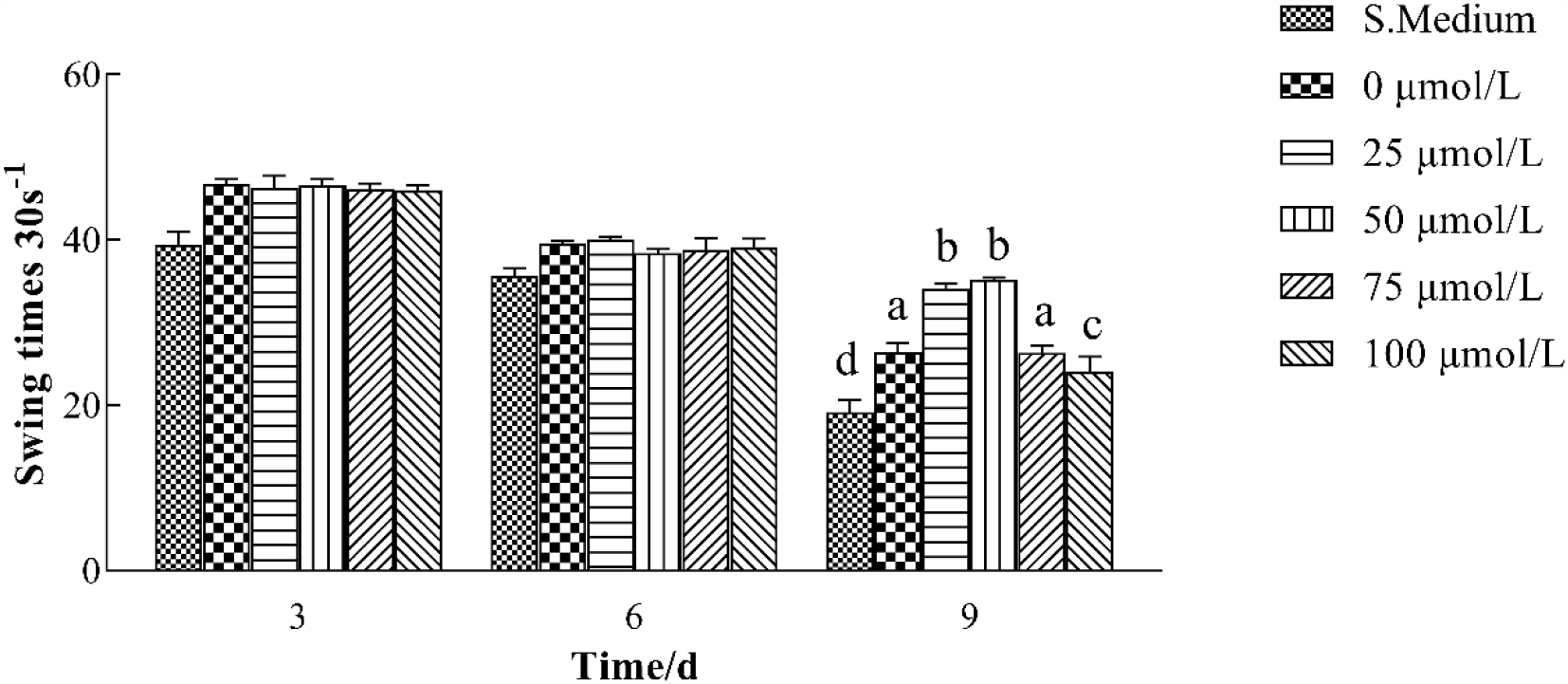
Effect of different concentrations of DHEA on swing times of *C. elegans* on day-3, day-6, day-9. Different letters represent significant differences between the groups of day-9.

### Effect of DHEA on intracellular ROS accumulation

The organism in the process of aging is often accompanied by the production of ROS. Over-accumulated ROS will react with DNA, protein, lipid in the body, causing oxidative damage to cells and tissues, accelerating the process of cell apoptosis and the organism’s aging. Diminishing the overproduction of intracellular ROS will benefit to extend the life-span of *C. elegans* (Labuschagne and Brenkman, 2013). As the result explained in Fig 5, the content of ROS in nematodes increased with time. The levels of intracellular ROS in worms exposed to DHEA (25 μmol/L, 50 μmol/L, 75 μmol/L and 100 μmol/L) were significantly lessened (P< 0.05). The finding fully proved that DHEA could decrease the quantity and delay the course of ROS accumulation in *C. elegans*, and then extend the life-span of *C. elegans*.

**Fig. 5.**
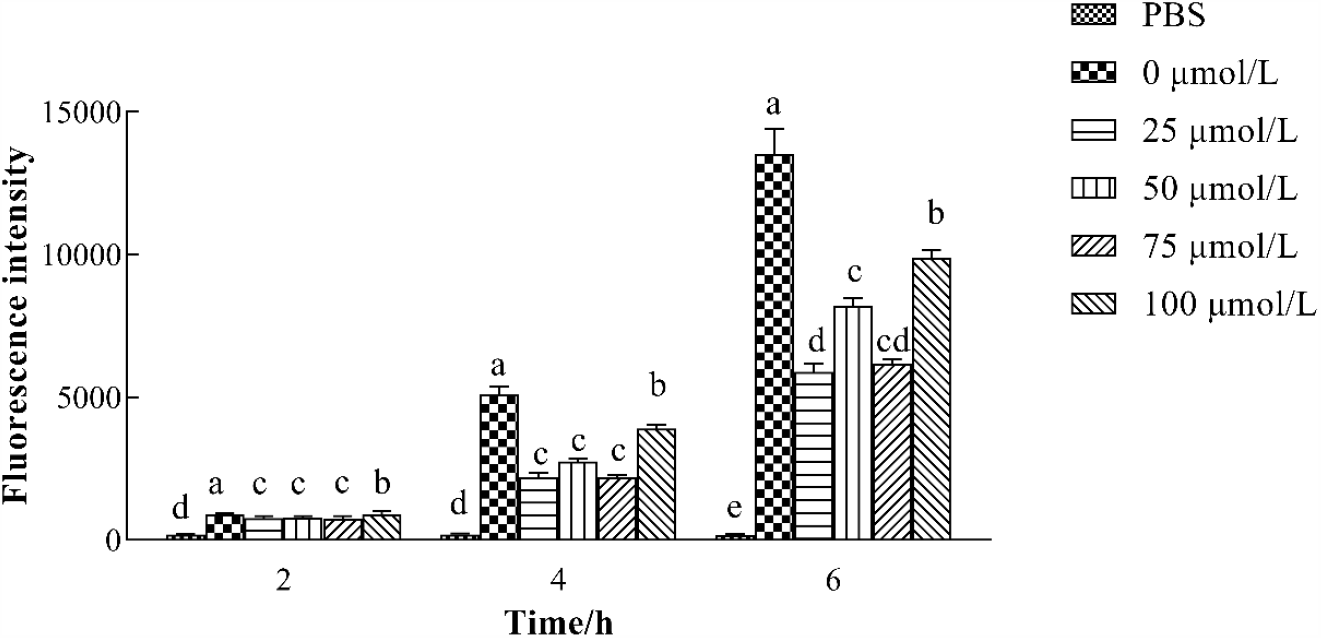
Mean(±SD) fluorescence intensity of *C. elegans* that treated with H_2_DCFDA. Different letters represent significant differences through 2^nd^, 4^th^ and 6^th^ h groups.

### Effect of DHEA on the content of lipofuscin in nematodes

Lipofuscin is an auto-fluorescent aging pigment, mainly composed of lipid peroxides and oxidized proteins. The amount of lipofuscin in most organisms including nematodes gradually increases with the age (Gerstbrein et al., 2005). The content of lipofuscin in *C. elegans* was detected and shown in Fig 6. Compared with the blank control group, the expression of lipofuscin significantly reduced in the worms cultivated with 50 μmol / L DHEA (P< 0.01), and no significant difference was found in the 25 μmol / L, 75 μmol / L DHEA and 25 μmol / L dose group and the S. Medium group (P> 0.05), while there was a significant increase in worms fed with 100 μmol / L DHEA (P<0.05).

**Fig. 6.**
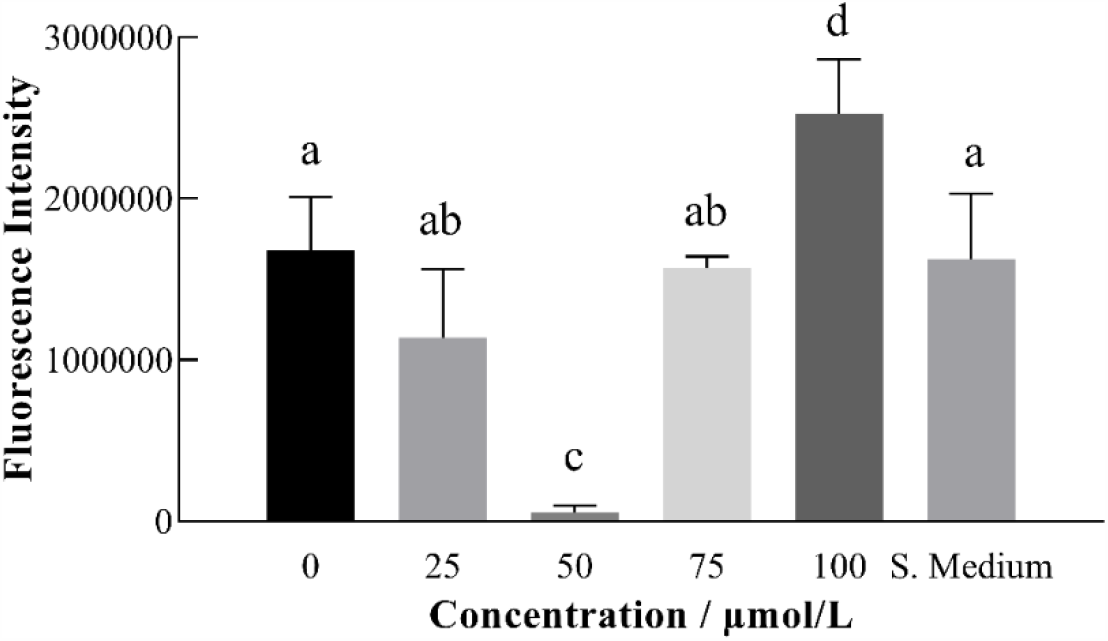
Effect of DHEA (0, 25, 50, 75, 100 μmol / L) on fluorescence intensity of *C. elegans*. Different letters indicate a significant difference between the 6 groups.

### Nuclear location of *daf-16*

The process of aging is influenced by two major aspects of genetics and environment. Currently accepted theories of aging include free radicals, telomeres and caloric restriction. *daf-16* gene is valuable to the longevity of nematodes, the nuclear expression of *daf-16* gene can significantly improve the antioxidant stress activity and the immunity of *C. elegans*, and leads a postponement of organisms’ aging (Zhang et al., 2013). To find out the antiaging mechanism of DHEA, TJ356 of transgenic mutant nematodes was taken for this study. Labeled the *daf-16* gene with green fluorescent protein (GFP), monitored the distribution of *daf-16* gene in cells with GFP. If DHEA acts on the Insulin-like growth factor (IGF) signaling pathway, the *daf-16* gene will be transferred from the cytoplasm into the nucleus to regulate the expression of a series of genes. Under the fluorescence microscope, spotted green fluorescence can be detected in nematode cell nucleus. Thus, we can judge whether DHEA regulates the nematode life through the IGF signal pathway. As shown in Fig 7, DHEA did distinctly promote the expression of *daf-16* in nucleus at the concentration of 50 μmol / L, and the fluorescence point in the nucleus appeared in this group only.

**Fig. 7.**
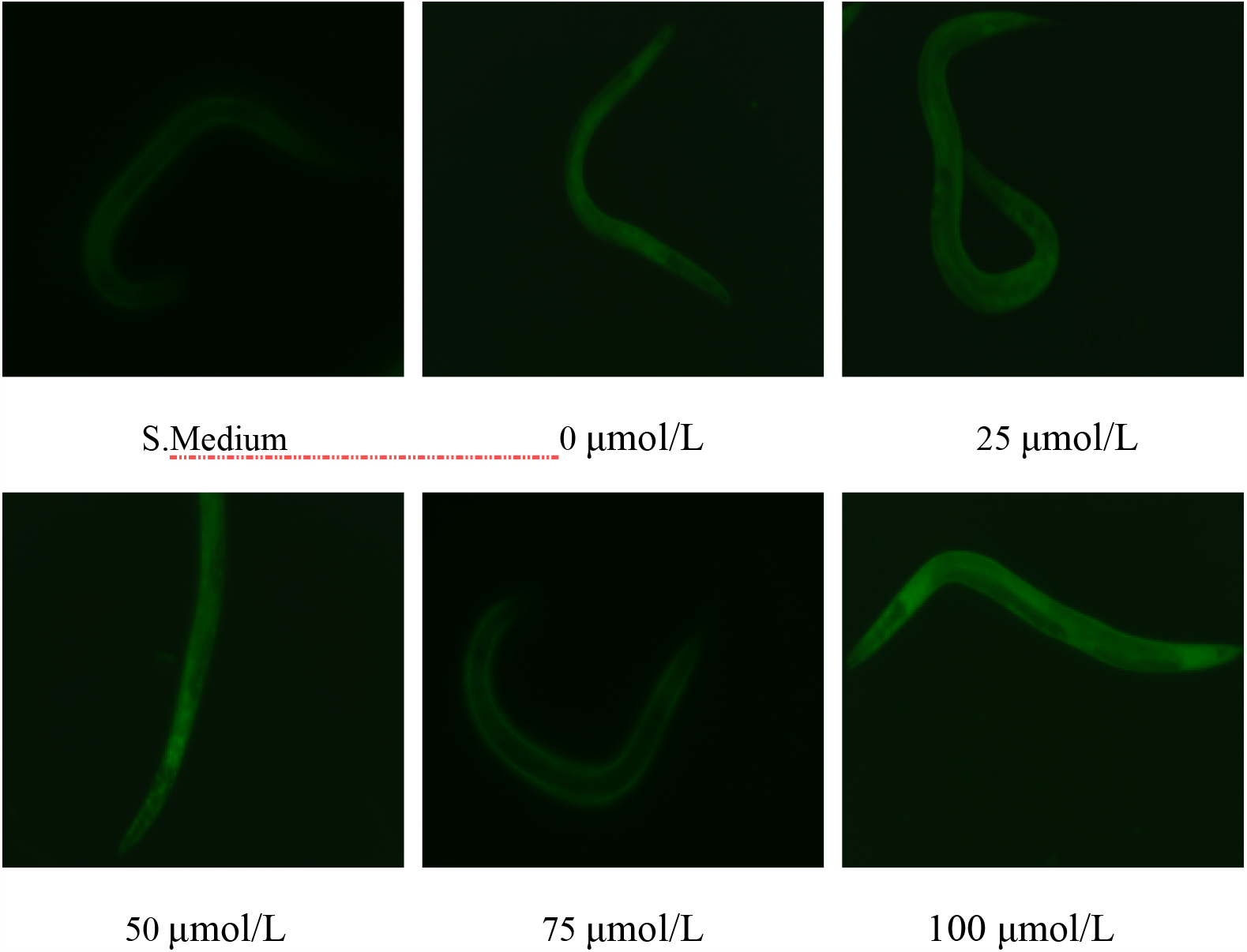
Distribution of fluorescent proteins in *C. elegans*.

## DISCUSSION

Current study was conducted to examine the effect of DHEA extract from sweet potato to articulate the life-extending ability using *Caenorhabditis elegans* model. The findings from this study articulated that DHEA was capable to increase the mean life-span and slowed down the growth velocity of body length and width, resisted the oxidative damage caused by H_2_O_2_, significantly reduced the content of ROS and lipofuscin of *C. elegans*. However, these effects not worked in a concentration-dependent manner. DHEA in 50 μmol / L could effectively enhance these abilities, while 100 μmol / L DHEA possesses a toxic effect on the growth of nematodes and could even accelerate aging process. Thus, we suspected that low concentration of DHEA had no significant effect on nematode senescence and there may exist a critical concentration between 50 μmol / L and 100 μmol / L, and beyond these levels, DHEA would accelerate the aging process of *C. elegans*. Further exploration suggested that the life extension of *C. elegans* supplied with DHEA was not accomplished through calorie restriction but in the way of promoting the nuclear expression of *daf-16* gene in *C. elegans*.

Although much is known about the beneficial effects of DHEA, there was few report explained the effect of DHEA on antiaging. The results from life-span assay indicated that DHEA may entered the nematodes’ body through the skin or esophagus to delay aging. In a parallel study, Wang et al. Added 400mg / ml of orange extract to nematode culture medium, and the mean life-span of nematodes was increased by 26.2%(Wang et al., 2020). And across table 2, we can see that DHEA would slow down the growth rate of body length and body width while delaying the life span. However, the extension of nematode life is determined by many factors, including insulin-like growth factor-1 signaling pathway, caloric restriction, stress resistance and oxidative damage (Chen et al., 2014b). Through the Fig 1 we can figure out that DHEA had no effect to the growth of *E. coli* OP_50_, and Fig 3 proved that DHEA would not affect the feeding behavior of *C. elegans*. However, it has been confirmed that calorie restrictions can greatly prolong the life-span of nematodes (Fang et al., 2016; Lakowski and Hekimi, 1998). So, we can draw the conclusion that the antiaging effect of DHEA on *C. elegans* was not realized through calorie restriction, which was coincided with the results of the mechanism of Rehmannia glutinosa neutral polysaccharide extending lifetime of *C. elegans* (Yuan et al., 2019). Moreover, the antiaging mechanism of DHEA on *C. elegans* needed further exploration.

In contrast, Fig 2 suggested that DHEA may delay the aging of *C. elegans* by improving their ability to resist external oxidative stress. The results were similar with Zhang’, which observed that the SOD activity, antioxidant stress and survival time of nematodes treated with *lycium barbarum* extract significantly increase(Zhang et al., 2020). However, the signaling pathways or functional factors that regulate the capacity of *C. elegans* to resist oxidative stress was still unknown.

In addition, Fig 4 and Fig 6 revealed that the effect of DHEA on the aging of *C. elegans* was not in a concentration-dependent manner. And there may be a critical concentration between 75 μmol / L and 100 μmol / L, which would change the effect of DHEA of life-span extension. The results here were similar with the finding of Panthakarn’s, which described the leaf extract of 50 μg/mL displayed the highest reduction of lipofuscin (Rangsinth et al., 2019). However, Song et al treated the worms with various concentrations of raspberry extract, the results showed that the extract can increase the motility of *C. elegans* in dose-dependent and time-dependent manner (Song et al., 2020). The finding fully proved that DHEA could decrease the quantity and delay the course of ROS accumulation in *C. elegans*, and then extend the life-span of *C. elegans*.

The results above proved that 50 μmol / L DHEA had a significant effect on the life-span, resistance to oxidative damage, and ROS and lipofuscin content of *C. elegans*. It is reported that *daf-16* gene encodes the FOXO / *daf-16* transcription factor in the insulin / IGF-1 signaling pathway, which directly affects the ability of *C. elegans* to resist stress(Lee et al., 2003; Murphy et al., 2003). The up-regulation of *daf-16* would enhance the expression of many effective downstream factors, such as superoxide dismutase, heat shock protein and choline transporter-like protein, so as to protect the *C. elegans* from stress and extend the longevity(Lin et al., 2002; Tissenbaum and Guarente, 2002). Through Fig 7, we guessed that these functions were realized by the regulation of *daf-16* gene in nematodes with the supply of DHEA. And the report from Cong also illustrated that fullerene facilitated the life-expanding effect of *C. elegans* by up-regulating the transcription level of *daf-16* gene (Cong et al., 2015).

In conclusion, these findings proved that the DHEA extract from sweet potato did have a favorable effect on life-extension. But further research about the effect of DHEA on *C. elegans* productive ability, resistance to thermotolerance stress, protein content, superoxide dismutase activity, catalase activity, glutathione enzyme activity and antiaging mechanism are needed to be carried out.

## Acknowledgments

We would like to thank Modassar Ranjha for his assistance with picture editing.

## Author contributions

HC drafted the manuscript, participated in the design of the study and analyzed the data; DZ carried out the experiments and participated in the design of the study; LFW provided the methodology and the funding support. All authors provided the final approval of the submission.

## References

Barkhausen, T., Hildebrand, F., Krettek, C. and van Griensven, M. (2009). DHEA-dependent and organ-specific regulation of TNF-alpha mRNA expression in a murine polymicrobial sepsis and trauma model. Critical Care 13.

Barrettconnor, E. and Edelstein, S. L. (1994). A PROSPECTIVE-STUDY OF DEHYDROEPIANDROSTERONE-SULFATE AND COGNITIVE FUNCTION IN AN OLDER POPULATION - THE RANCHO-BERNARDO STUDY. Journal of the American Geriatrics Society 42, 420–423.

Chege, P. M. and McColl, G. (2014). Caenorhabditis elegans: a model to investigate oxidative stress and metal dyshomeostasis in Parkinson’s disease. Frontiers in Aging Neuroscience 6.

Chen, W.-C., Hsieh, S.-R., Chiu, C.-H., Hsu, B.-D. and Liou, Y.-M. (2014a). Molecular identification for epigallocatechin-3-gallate-mediated antioxidant intervention on the H2O2-induced oxidative stress in H9c2 rat cardiomyoblasts. J Biomed Sci 21.

Chen, Y., Onken, B., Chen, H., Xiao, S., Liu, X., Driscoll, M., Cao, Y. and Huang, Q. (2014b). Mechanism of longevity extension of Caenorhabditis elegans induced by pentagalloyl glucose isolated from eucalyptus leaves. J Agric Food Chem 62, 3422–31.

Cong, W., Wang, P., Qu, Y., Tang, J., Bai, R., Zhao, Y., Chunying, C. and Bi, X. (2015). Evaluation of the influence of fullerenol on aging and stress resistance using Caenorhabditis elegans. Biomaterials 42, 78–86.

Deleon, M. J., Horani, M. H., Haas, M. J., Wong, N. C. W. and Mooradian, A. D. (2002). Effects of dehydroepiandrosterone on rat apolipoprotein Al gene expression in the human hepatoma cell line, HepG2. Metabolism-Clinical and Experimental 51, 376–379.

Fang, B., Zhang, M., Ren, F. Z. and Zhou, X. D. (2016). Lifelong diet including common unsaturated fatty acids extends the lifespan and affects oxidation in Caenorhabditis elegans consistently with hormesis model. European Journal of Lipid Science and Technology 118, 1084–1092.

Genazzani, A. D., Lanzoni, C. and Genazzani, A. R. (2007). Might DHEA be considered a beneficial replacement therapy in the elderly? Drugs & Aging 24, 173–185.

Gerstbrein, B., Stamatas, G., Kollias, N. and Driscoll, M. (2005). In vivo spectrofluorimetry reveals endogenous biomarkers that report healthspan and dietary restriction in Caenorhabditis elegans. Aging Cell 4, 127–137.

Grillon, C., Pine, D. S., Baas, J. M. P., Lawley, M., Ellis, V. and Charney, D. S. (2006). Cortisol and DHEA-S are associated with startle potentiation during aversive conditioning in humans. Psychopharmacology (Berl) 186, 434–441.

Ku, H.-C., Chen, W.-P. and Su, M.-J. (2013). DPP4 Deficiency Exerts Protective Effect against H2O2 Induced Oxidative Stress in Isolated Cardiomyocytes. PLoS One 8.

Labuschagne, C. F. and Brenkman, A. B. (2013). Current methods in quantifying ROS and oxidative damage in Caenorhabditis elegans and other model organism of aging. Ageing Research Reviews 12, 918–930.

Lakowski, B. and Hekimi, S. (1998). The genetics of caloric restriction in Caenorhabditis elegans. Proceedings of the National Academy of Sciences of the United States of America 95, 13091–13096.

Lee, S. S., Kennedy, S., Tolonen, A. C. and Ruvkun, G. (2003). DAF-16 target genes that control C-elegans life-span and metabolism. Science 300, 644–647.

Lin, S. J., Kaeberlein, M., Andalis, A. A., Sturtz, L. A., Defossez, P. A., Culotta, V. C., Fink, G. R. and Guarente, L. (2002). Calorie restriction extends Saccharomyces cerevisiae lifespan by increasing respiration. Nature 418, 344–348.

Loening-Baucke, V., Miele, E. and Staiano, A. (2004). Fiber (glucomannan) is beneficial in the treatment of childhood constipation. Pediatrics 113, E259–E264.

Ludewig, A. H., Kober-Eisermann, C., Weitzel, C., Bethke, A., Neubert, K., Gerisch, B., Hutter, H. and Antebi, A. (2004). A novel nuclear receptor/coregulator complex controls C. elegans lipid metabolism, larval development, and aging. Genes & Development 18, 2120–2133.

Masoro, E. J. (2005). Overview of caloric restriction and ageing. Mech Ageing Dev 126, 913–922.

Mohanraj, R. and Sivasankar, S. (2014). Sweet potato (Ipomoea batatas [L.] Lam)--a valuable medicinal food: a review. J Med Food 17, 733–41.

Murphy, C. T., McCarroll, S. A., Bargmann, C. I., Fraser, A., Kamath, R. S., Ahringer, J., Li, H. and Kenyon, C. (2003). Genes that act downstream of DAF-16 to influence the lifespan of Caenorhabditis elegans. Nature 424, 277–284.

Nordmark, G., Bengtsson, C., Larsson, A., Karlsson, F. A., Sturfelt, G. and Rnnblom, L. (2005). Effects of dehydroepiandrosterone supplement on health-related quality of life in glucocorticoid treated female patients with systemic lupus erythematosus. Autoimmunity 38, 531–540.

Ogawa, T., Kodera, Y., Hirata, D., Blackwell, T. K. and Mizunuma, M. (2016). Natural thioallyl compounds increase oxidative stress resistance and lifespan in Caenorhabditis elegans by modulating SKN-1/Nrf. Sci Rep 6.

Pant, A., Saikia, S. K., Shukla, V., Asthana, J., Akhoon, B. A. and Pandey, R. (2014). Beta-caryophyllene modulates expression of stress response genes and mediates longevity in Caenorhabditis elegans. Experimental Gerontology 57, 81–95.

Peixoto, H., Roxo, M., Silva, E., Valente, K., Braun, M., Wang, X. and Wink, M. (2019). Bark Extract of the Amazonian Tree Endopleura uchi (Humiriaceae) Extends Lifespan and Enhances Stress Resistance in Caenorhabditis elegans. Molecules 24.

Ran, J., Liang, X., Du, H. and Sun, J. (2019). Optimization of DHEA Extraction from Sweet Potato Pomace by Ultrasonic-Microwave Synergistic Employing Response Surface Methodology. J AOAC Int 102, 680–682.

Rangsinth, P., Prasansuklab, A., Duangjan, C., Gu, X., Meemon, K., Wink, M. and Tencomnao, T. (2019). Leaf extract of Caesalpinia mimosoides enhances oxidative stress resistance and prolongs lifespan in Caenorhabditis elegans. BMC Complement Altern Med 19, 164.

Ruan, Q., Qiao, Y., Zhao, Y., Xu, Y., Wang, M., Duan, J. and Wang, D. (2016). Beneficial effects of Glycyrrhizae radix extract in preventing oxidative damage and extending the lifespan of Caenorhabditis elegans. Journal of Ethnopharmacology 177, 101–110.

Sohal, R. S., Ku, H. H., Agarwal, S., Forster, M. J. and Lal, H. (1994). OXIDATIVE DAMAGE, MITOCHONDRIAL OXIDANT GENERATION AND ANTIOXIDANT DEFENSES DURING AGING AND IN RESPONSE TO FOOD RESTRICTION IN THE MOUSE. Mech Ageing Dev 74, 121–133.

Song, B., Zheng, B., Li, T. and Liu, R. H. (2020). Raspberry extract promoted longevity and stress tolerance via the insulin/IGF signaling pathway and DAF-16 in Caenorhabditis elegans. Food Funct.

Tissenbaum, H. A. and Guarente, L. (2002). Model organisms as a guide to mammalian aging. Developmental Cell 2, 9–19.

Viswanathan, M., Kim, S. K., Berdichevsky, A. and Guarente, L. (2005). A role for SIR-2.1 regulation of ER stress response genes in determining C-elegans life span. Developmental Cell 9, 605–615.

Wang, D. and Xing, X. (2010a). Pre-treatment with mild UV irradiation suppresses reproductive toxicity induced by subsequent cadmium exposure in nematodes. Ecotoxicology and Environmental Safety 73, 423–429.

Wang, D. Y. and Xing, X. J. (2010b). Pre-treatment with mild UV irradiation suppresses reproductive toxicity induced by subsequent cadmium exposure in nematodes. Ecotoxicology and Environmental Safety 73, 423–429.

Wang, J., Deng, N., Wang, H., Li, T., Chen, L., Zheng, B. and Liu, R. H. (2020). Effects of Orange Extracts on Longevity, Healthspan, and Stress Resistance in Caenorhabditis elegans. Molecules 25.

Webb, S. J., Geoghegan, T. E. and Prough, R. A. (2006). The biological actions of dehydroepiandrosterone involves multiple receptors. Drug Metabolism Reviews 38, 89–116.

Yuan, Y., Kang, N., Li, Q., Zhang, Y., Liu, Y. and Tan, P. (2019). Study of the Effect of Neutral Polysaccharides from Rehmannia glutinosa on Lifespan of Caenorhabditis elegans. Molecules 24.

Zhang, J., Ji, D.-S., Zhao, S.-H., Xie, X.-L., Chen, H. and Li, H.-F. (2020). Anti-Aging Effects of Lycium ruthenicum Murr. Granules in Caenorhabditis elegans. Pakistan Journal of Zoology 52.

Zhang, P., Judy, M., Lee, S.-J. and Kenyon, C. (2013). Direct and Indirect Gene Regulation by a Life-Extending FOXO Protein in C. elegans: Roles for GATA Factors and Lipid Gene Regulators. Cell Metabolism 17, 85–100.

